# Nuclear exclusion of menin drives functional MEN1 deficiency in non-MEN1 prolactinomas: mouse models and human biopsies

**DOI:** 10.64898/2026.08.19.745771

**Authors:** M Peña-Zanoni, Á Flores-Martínez, D Bornancini, AI Abeledo Machado, Victoria Segobia, SB Rulli, RM Luque, G Díaz-Torga

**Author notes:** **Corresponding author to whom reprint requests should be addressed:** Graciela Díaz-Torga Instituto de Biología y Medicina Experimental-CONICET Vuelta de Obligado 2490, Buenos Aires 1428, Argentina. Equal contribution (RM Luque, G Díaz-Torga).

## Abstract

Prolactinomas, the most common secretory pituitary tumour subtype, frequently occur in patients with Multiple Endocrine Neoplasia type 1, caused by germline MEN1 mutations encoding menin. While menin loss is well established in MEN1-associated prolactinomas, its role in sporadic tumours remains unclear. We investigated menin expression, subcellular localization, and downstream signalling in two murine models of non-MEN1 prolactinomas, the dopamine D2-receptor knockout and the hCGβ-subunit-overexpressing mice, in which only females develop prolactinoma. Pituitary *Men1* expression, analysed by qPCR, remained unchanged despite the genotype, in both sexes. However, in prolactinomas, lactotrophs exhibited a marked loss of nuclear MEN1 immunostained, with protein restricted to the cytoplasm. Male mice pituitaries retained nuclear MEN1 localization regardless their genotype. Loss of nuclear menin in prolactinomas was associated with reduced *p27* and *Pten* expression, increased *Ccnd1* expression, and enhanced pAKT. Moreover, by using *in vivo* pharmacological and surgical approaches we demonstrated that dopamine-agonist treatment preserved nuclear menin in lactotrophs, whereas dopamine blockade or estradiol induced its nuclear loss. Importantly, analysis of human pituitary biopsies confirmed nuclear and cytoplasmic menin localization in lactotrophs from normal pituitaries, and in prolactinomas from both genders following dopamine agonist therapy. However, in a prolactinoma from an untreated female, nuclear menin was partially lost. Therefore, our findings identify a state of functional MEN1-deficiency in sporadic prolactinomas (characterized by preserved MEN1 expression), but its exclusion from the nucleus (linked to activation of proliferative pathways, impaired tumour suppressor signalling, and tumour development) highlights the restoration of nuclear MEN1 localization as a potential therapeutic strategy.

**Graphical abstract:** Novel mechanism of functional MEN1 deficiency

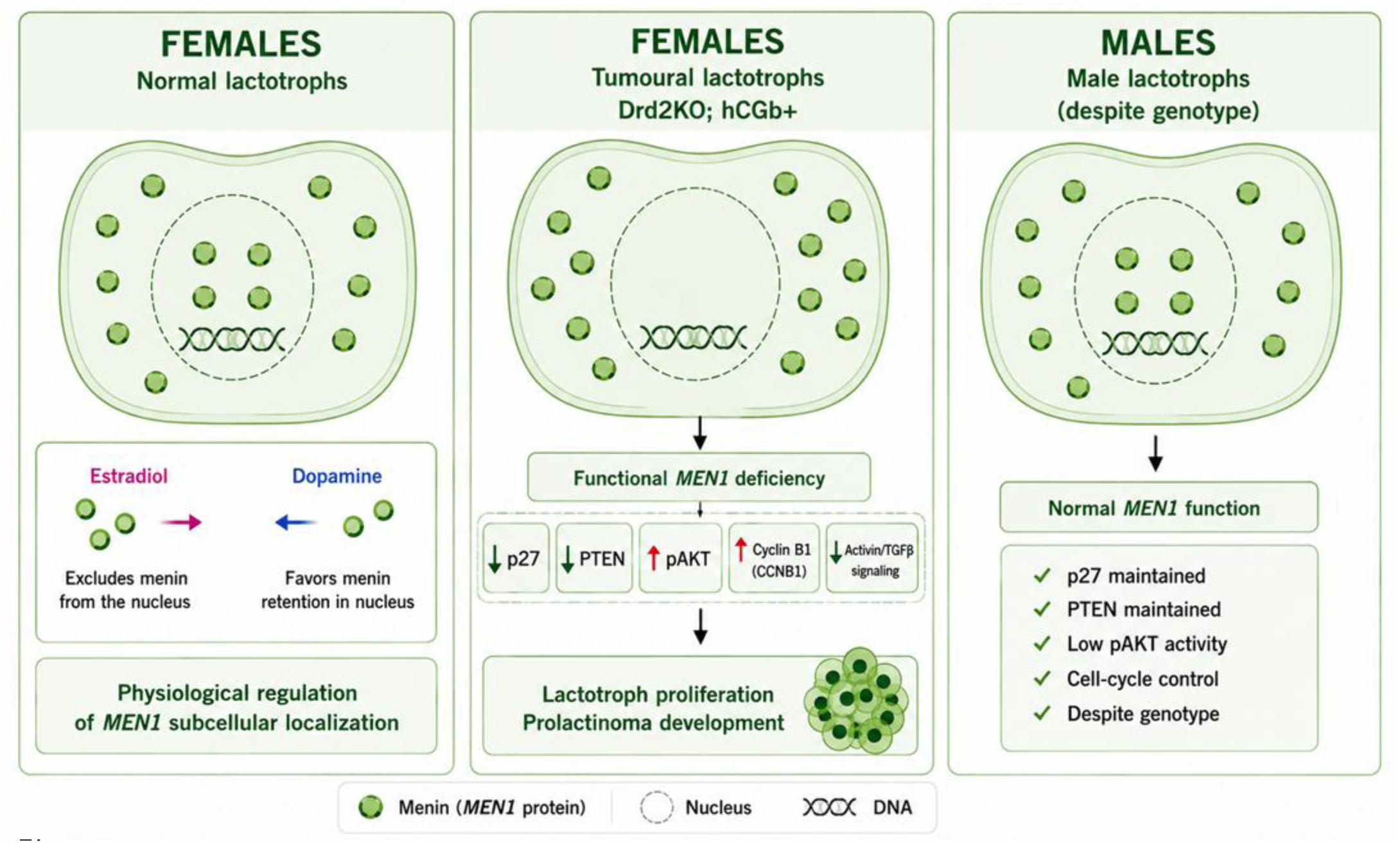

## INTRODUCTION

Menin was first identified in the late 1990s through genetic linkage analysis as the causative gene of Multiple Endocrine Neoplasia type 1 (MEN1), a rare hereditary syndrome caused by germline mutations in the *MEN1* gene located on chromosome 11q13. It follows an autosomal dominant inheritance pattern with high penetrance, characterised by the development of multiple endocrine tumours, primarily affecting the parathyroid glands, pancreas, and anterior pituitary, often presenting in early adulthood (1)(2). Menin plays a key role in cell cycle regulation, genome stability, DNA repair, and cell proliferation by influencing the activity and/or expression of several regulators, either through direct interaction or by modulating the transcription of their genes (3,4). It functions as a tumour suppressor in endocrine organs by blocking the G0/G1 to S phase transition. This occurs through the downregulation of cyclin-dependent kinase 2 (CDK2) expression and activity, and the upregulation of CDKIs such as *p27^Kip1^* and *p18^Ink4c^* (5)(6)(7). Additionally, menin interacts with AKT to downregulate pAKT-induced proliferation in both endocrine and non-endocrine cells (8).

Menin also acts as a scaffold protein, interacting with a broad range of partners, including transcription factors such as JunD and nuclear receptors, and members of the mixed lineage leukaemia histone methyltransferase complex, exerting both repressive and activating effects on target genes involved in cell cycle control and differentiation, depending on the cell type (9)(10)(11).

Loss-of-function mutations in *MEN1* disrupt these regulatory pathways, resulting in dysregulated gene transcription and increased cellular proliferation, which underlie the tumorigenic process in MEN1-associated tissues. Interestingly, although menin is broadly expressed in many tissues, tumour formation in MEN1 patients occurs predominantly in endocrine organs, highlighting the existence of tissue-specific regulatory roles and molecular interactions that remain poorly understood.

Prolactinomas represent the most prevalent pituitary tumour subtype observed in MEN1 patients (12). Although MEN1 mutations are well-established drivers of hereditary prolactinomas, the role of menin in sporadic prolactinomas remains poorly defined (13)(14). Most studies have focused on MEN1 gene expression levels, with conflicting results regarding its contribution to pituitary tumorigenesis. Importantly, menin function depends not only on its expression but also on its subcellular localization, as many of its tumour suppressor activities require nuclear residency. Therefore, we hypothesized that alterations in cellular menin localization rather than MEN1 expression may contribute to prolactinoma development. To test this hypothesis, we investigated MEN1 expression, intracellular localization, and downstream signalling pathways in two independent mouse models of non-MEN1 prolactinomas and explored the contribution of dopamine and estradiol to these processes. In addition, we presented a preliminary study of menin subcellular localization in lactotrophs from human pituitary biopsies.

## MATERIALS AND METHODS

### Animal models

The *Drd2* knock-out mice model (official strain designation B6; 129S2-*Drd2^tm1low^*: JAX stock #003190) was generated by targeted mutagenesis of the dopamine receptor type 2 gene (*Drd2*) in embryonic stem cells (15)(16). These mice lack functional dopamine receptors type 2, neither the long nor the short isoform. These mice were used at the age of eight months, age at which Drd2ko females show pronounced hyperplastic pituitaries and hyperprolactinemia.

The hCGβ+ mice model, with FVB/N background, overexpresses the human chorionic gonadotrophin β (hCGβ) subunit under the control of ubiquitin C promoter. hCGβ+ female mice develop precocious puberty, have increased levels of progesterone and testosterone, present infertility with severe disturbances in their reproductive system, and develop mammary tumours and prolactinomas (17). These mice were used at age of six months. At this time females present a rise in PRL secretion and their pituitaries show pronounced lactotroph hypertrophy.

In both models, wild-type (WT) littermates were used as controls. Mice were housed in a temperature-controlled room under a 12:12 h light-dark cycle, with free access to laboratory chow and filtered tap water. The experimental protocols followed the NIH Guidelines for Care and Use of Experimental Animals (Division of Animal Welfare, Office for Protection of Research Risks, National Institutes of Health, Assurance identification number F16-00065, A#5072-01). All experimental protocols were also approved by the Institutional Animal Care and Use Committee of the “Instituto de Biología y Medicina Experimental” from the “Consejo Nacional de Investigaciones Científicas y Técnicas” (IBYME-CONICET), number 019/2024.

After euthanasia by decapitation, pituitaries were removed, without the neural lobe, and stored in Quick-Zol (Kalium technologies) at-70 °C for subsequent RNA isolation and gene expression studies by real-time PCR, or Cryoplast (Biopack) at −70 °C for subsequent protein analysis by immunofluorescence detection.

### *In vivo* experiments

Wild-type female mice (Drd2 model) received intraperitoneal (i.p.) injection of either saline solution (control group), the Drd2 antagonist sulpiride (10 mg/kg; IVAX Laboratories), or the Drd2 agonist cabergoline (2 mg/kg; Beta Laboratories, Buenos Aires). Another group received subcutaneous injection of estradiol-valerate (0.2 mg/kg, Progynon® Depot, Schering) or castor oil (control group). Animals were euthanized by decapitation after 3 hours of treatment, and anterior pituitaries were processed for immunostaining, as described below.

### Ovariectomy (OVX), the surgical procedure

Female hCGβ+ transgenic mice underwent bilateral oophorectomy or sham surgery at two months of age under ketamine/xylazine anaesthesia (90 mg/kg and 10 mg/kg, respectively; i.p.). At six months of age, animals were euthanized by decapitation, and the anterior pituitaries were collected and processed for immunostaining as described below.

### RNA isolation and Real Time PCR

Anterior pituitaries, collected in Quick-Zol (Kalium technologies), were processed for RNA isolation following the manufacturer’s protocol. Briefly, the samples were homogenized, mixed with chloroform for nucleic acid phase separation and then mixed with isopropanol for RNA precipitation. For each sample, one microgram of RNA was reverse-transcribed using the MMLV-RT enzyme (Promega) and random primers (Biodynamics). The cDNA obtained was used for gene expression analysis. Primers sets were designed for the specific amplification of murine *Men1*, *Ccnd1*, *Ccne1*, *Pten*, *Rpl38* and *CyclophilinB*. Table 1 shows the primer sequences used.

**Table 1:**
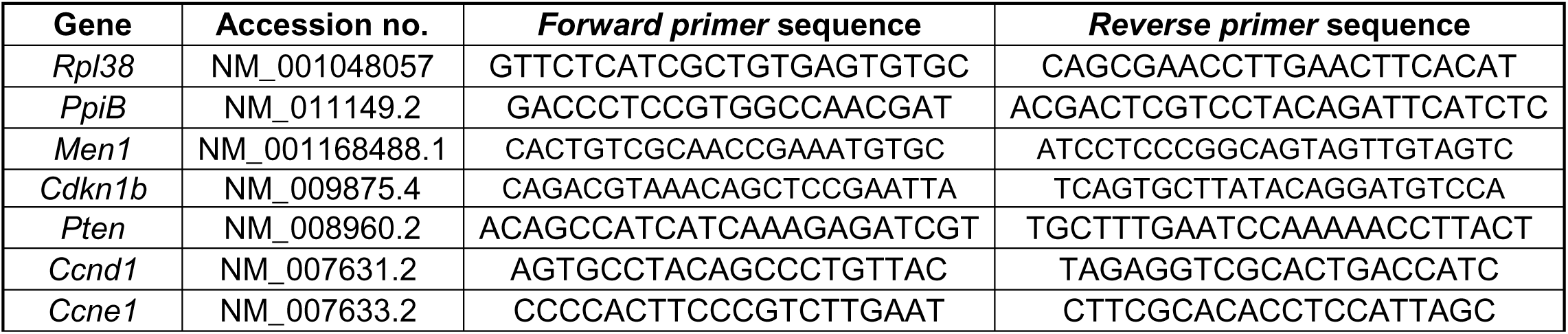
Real Time PCR primer sequences.

For reverse-transcription quantitative PCR (RT-qPCR) assays, the Fast Start Universal SYBR Green Master Rox (Roche) or the HOT FIREPol® EvaGreen® qPCR Mix Plus (Solis BioDyne) were used with the respective temperature and time protocols, on a CFX96 Touch Real-Time PCR Detection System (BioRad). The most appropriate reference gene for each gene of interest was selected by performing qPCR efficiency studies with serially diluted samples and two reference genes (*CyclophilinB* and *Rpl38*). The threshold cycle (Ct) of the target gene was normalized to the corresponding reference gene, and the mean value of the WT female group was used as the calibrator. Differences in gene expression were quantified using the Pfaffl method (18), where relative expression ratios were calculated by correcting for differences in primer amplification efficiencies.

### Double indirect immunostaining in mouse pituitaries

Pituitaries placed in Cryoplast (Biopack) at-70°C were sectioned using a cryostat at-20°C and the immunofluorescence technique was performed as previously described (19). Briefly, tissue sections were fixed in absolute methanol, then blocked with 5% PBS–BSA for 1 h and finally incubated overnight at 4°C with the corresponding primary antibody: anti-menin (1:50; sc-374371, Santa Cruz Biotechnology), anti-p27 (1:50; sc-1641, Santa Cruz Biotechnology) or anti-pAKT (1:100; sc-7985, Santa Cruz Biotechnology) (Table 2).

**Table 2:** Primary and secondary antibodies used in immunofluorescence assays.

| Primary antibodies |  |  |  |  |  |
| --- | --- | --- | --- | --- | --- |
| Antibody | Brand | Catalog N° | Isotype | Specie | Dilution |
| αPRL | Parlow | AFP65191 | IgG | Guinea Pig | 1:1000 |
| αMenin | Santa Cruz | sc-374371 | IgG | Mouse Monoclonal | 1:50 |
| αp27 | Santa Cruz | sc-1641 | IgG | Mouse Monoclonal | 1:50 |
| αpAKT | Santa Cruz | sc-7985 | IgG | Rabbit Polyclonal | 1:100 |
| αPRL | Roche | 5268249001 | IgG | Rabbit Polyclonal | 1:25 |
| Secondary antibodies |  |  |  |  |  |
| Antibody | Brand | Catalog N° | Fluorescent dye |  | Dilution |
| αGuineaPig IgG | Abcam | ab150185 | Alexa 488 |  | 1:1000 |
| αMouse IgG | Santa Cruz | sc-516179 | CFL 555 |  | 1:200 |
| αRabbit IgG | Santa Cruz | sc-516251 | CFL 647 |  | 1:200 |
| αMouse IgG | Invitrogen | A-21043 | Alexa 488 |  | 1:1000 |
| αRabbit IgG | Invitrogen | A-78955 | Alexa 568 |  | 1:1000 |

The next day, the sections were incubated with anti-PRL (1:1000; AFP65191; Dr A. Parlow, National Hormone and Pituitary Program, Torrance, CA, USA) for 2 h at room temperature. After that, the sections were incubated with secondary antibodies conjugated with CFL-647 (dilution 1:200; sc-516251, Santa Cruz Biotechnology) or CFL-555 (1:100; sc-516179, Santa Cruz Biotechnology), and Alexa 488 (dilution 1:1000; ab150185, Abcam) (Table 2) for 1.5 h at room temperature in darkness, in a wet chamber. Between the mentioned steps, the sections were washed with PBS 1x. Finally, tissues were incubated for fifteen minutes with 4′,6-diamidino-2-phenylindole (DAPI) and then mounted with Fluoromount (Sigma, St Louis, MO). To validate the specificity of each immunostaining, tissues incubated with 1% PBS–BSA instead of the primary antibody were used as negative controls.

Images at 60x were obtained from an inverted confocal laser-scanning microscope Olympus IX83 with Spinning Disk Unit (Olympus; Tokyo, Japan) or the confocal microscope Olympus FV10i. Images were analysed with the Fiji software. For each experimental group, three or four slides, corresponding to different pituitaries, were analysed. A total of 10000 pituitary cells, in randomly chosen fields of each glass slide, was quantified, and the percentage of lactotroph cells (PRL+) expressing menin, p27 or pAKT was established.

### Immunofluorescence analysis of human biopsies

Formalin-fixed paraffin-embedded (FFPE) human pituitary samples, including prolactinomas and non-tumoral control pituitary tissues, were obtained through the Andalusian Public Health System Biobank (Córdoba Node; Spain). The Biobank coordinated the collection, processing, management and assignment of the biological samples used in this study according to its standardized procedures. Control pituitary tissues were obtained from autopsy and craniotomy specimens, as previously described (20). Clinicopathological information for the human samples included in the study is provided in Table 3. All samples were histologically evaluated by expert pathologists to confirm the diagnosis and, in the case of tumour specimens, to ensure the presence of representative tumour tissue in the analysed sections.

**Table 3:** Clinicopathological information for the human samples for immunofluorescence protocols.

| Tumor | Age at surgery (years) | Gender | Diagnosis (adenoma) | Cabergoline pretreatment | Other presurgery treatment |
| --- | --- | --- | --- | --- | --- |
| PRLoma#01 | 37 | Male | Yes | Yes |  |
| PRLoma#02 | 21 | Female | Yes | Yes | Eutirox 25 mg; hydrocortisone 20 mg; and diane 35 mg |
| PRLoma#03 | 24 | Female | Yes | No | Levonorgestrel/ethinylestradiol; losartan 50 mg; atorvastatin 20 mg; folic acid; desloratadine; and furosemide |

| Normal pituitaries | Age (years) | Gender | Peritumoral | Autopsy |
| --- | --- | --- | --- | --- |
| NP#01 | 66 | Male | No | Yes |
| NP#02 | 40 | Female | No | Yes |

All procedures involving human samples were conducted in accordance with the ethical principles of the Declaration of Helsinki of the World Medical Association. The study was approved by the corresponding Ethics Committee (Comité Coordinador de Ética de la Investigación Biomédica de Andalucía, approval code: S2500301). Written informed consent was obtained from all individuals included in the study or, when applicable, from a family member or legally authorized representative.

Immunofluorescence analysis was performed as previously described (21). Briefly, 5 μm-thick FFPE tissue sections were deparaffinized, rehydrated through xylene, graded ethanol solutions, and washed in distilled water. Heat-induced epitope retrieval was performed using Tris–EDTA buffer, pH 9.0. Sections were then permeabilized with 0.2% Triton X-100 in PBS and blocked for 45 min at room temperature (RT) using blocking solution containing 10% normal goat serum and BSA in PBS. Sections were sequentially incubated with the following primary antibodies: mouse anti-menin1 antibody (1:25; sc-374371, Santa Cruz Biotechnology) and anti-prolactin antibody (1:25; 05268249001, VENTANA, Roche Diagnostics) (Table 2). After washing with PBS, sections were incubated for 45 min at RT, protected from light, with the corresponding species-specific secondary antibodies conjugated to Alexa Fluor 488 or Alexa Fluor 568 (Thermo Fisher Scientific). Nuclei were counterstained with DAPI and slides were mounted using a fluorescence-compatible mounting medium (DAKO, S302380-2). Negative controls were performed by replacing the primary antibodies with blocking solution. Immunofluorescence images were acquired using a ZEISS LSM-7 DUO confocal system (Carl Zeiss, Oberkochen, Germany) maintaining comparable acquisition settings across samples.

### Statistical analysis

Student’s t-test was used to compare the results from two groups (gene expression or protein expression values). Two-way ANOVA was performed when analysing the effects of two factors: genotype and sex, followed by Tukey’s post hoc test when the interaction effect was significant: p<0.05. One-way ANOVA was performed to compare the means of three independent groups (WT vs hCGβ+ vs hCGβ+OVX analyses), followed by Bonferroni’s *post hoc* test when significant interaction. The analysis was performed using the GraphPad Prism 9 software. Results were expressed as mean ± SEM, and the p values are detailed in each figure legend.

## RESULTS

### *Men1* mRNA expression is not altered in prolactinomas from Drd2 and hCGβ mice

We quantified *Men1* mRNA expression in pituitaries from both mice models, including both sexes and genotypes (Figure 1). We observed no differences in pituitary *Men1* expression between sexes or genotypes in the Drd2 model (Fig. 1A). However, in the hCGβ+ model (Fig. 1B), male mice exhibited higher pituitary *Men1* expression levels than females, with no significant differences between genotypes. Overall, these findings indicate that *Men1* gene expression is not altered in prolactinomas compared to WT pituitaries.

**Figure 1:**
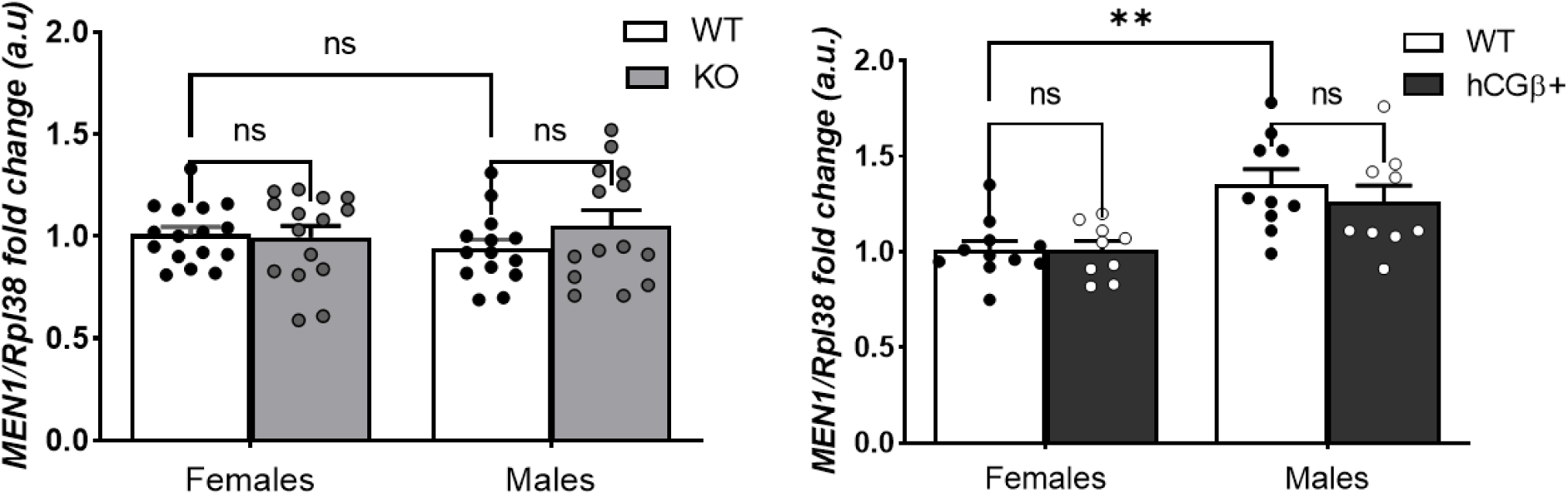
Pituitary mRNA expression of MEN1 in Drd2 and hCGβ mouse models. mRNA transcripts were amplified with specific primers by qRT-PCR and normalized to *Rpl38*. Results are expressed relative to those for WT females. (A) 8-month-old Drd2 mice. Interaction ns. (B) 6-month-old hCGβ mice. Interaction ns. \*\**p* = < 0.01 WT male vs WT female (sex difference). ns = not significant. Data were analysed by two-way ANOVA (sex x genotype). Data are expressed as mean ± SEM. n=9-16/group.

### Changes in MEN1 subcellular localization in prolactinomas from Drd2KO and hCGβ+ female mice

We performed double indirect immunofluorescence to evaluate menin protein expression specifically in lactotrophs. We then analyzed confocal microscopy images and quantified cells immunoreactive for both PRL and menin (PRL⁺/Men⁺) (Figure 2, Figure 3). We observed no significant differences in the percentage of PRL⁺/Men⁺ cells when comparing prolactinomas (pituitaries from Drd2KO or hCGβ⁺ females) with their respective WTs females (Fig. 2C and 3C), nor between transgenic males and their WT counterparts (Fig. 2D and 3D).

**Figure 2:**
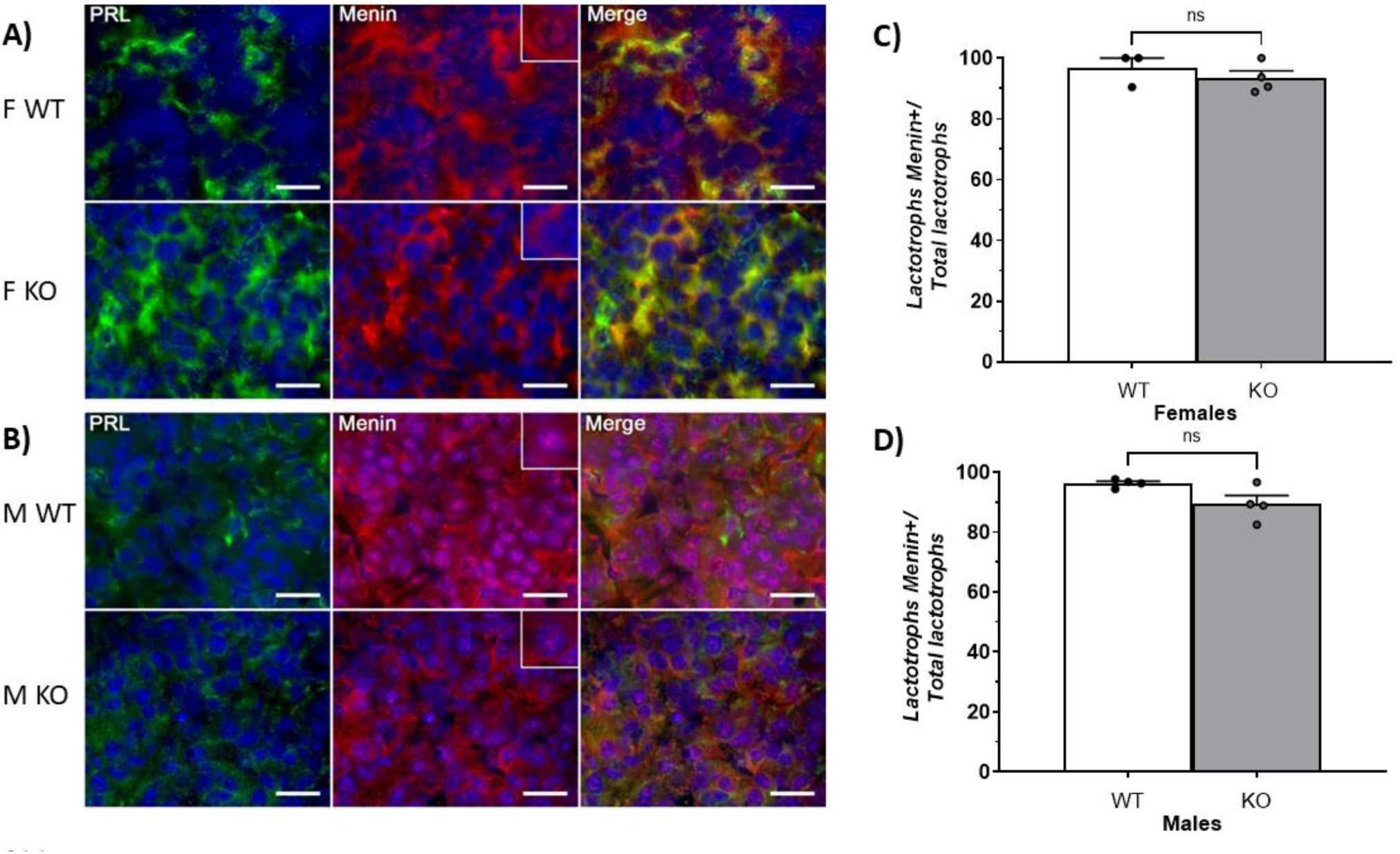
Menin protein expression in lactotrophs from Drd2 mice pituitaries. (A-B) Immunodetection of menin (red) and prolactin (green) by double indirect immunofluorescence in WT and KO female (A) and male (B) mice pituitaries. Nuclei were immunostained with DAPI (blue). Scale bar = 20 μm. **(C-D)** Percentage of lactotrophs expressing Menin/total lactotrophs in female (C) and male (D) mice pituitaries. ns = not significant. Data analysed by Student t test. Data are expressed as mean ± SEM. n = 3-4/group.

**Figure 3:**
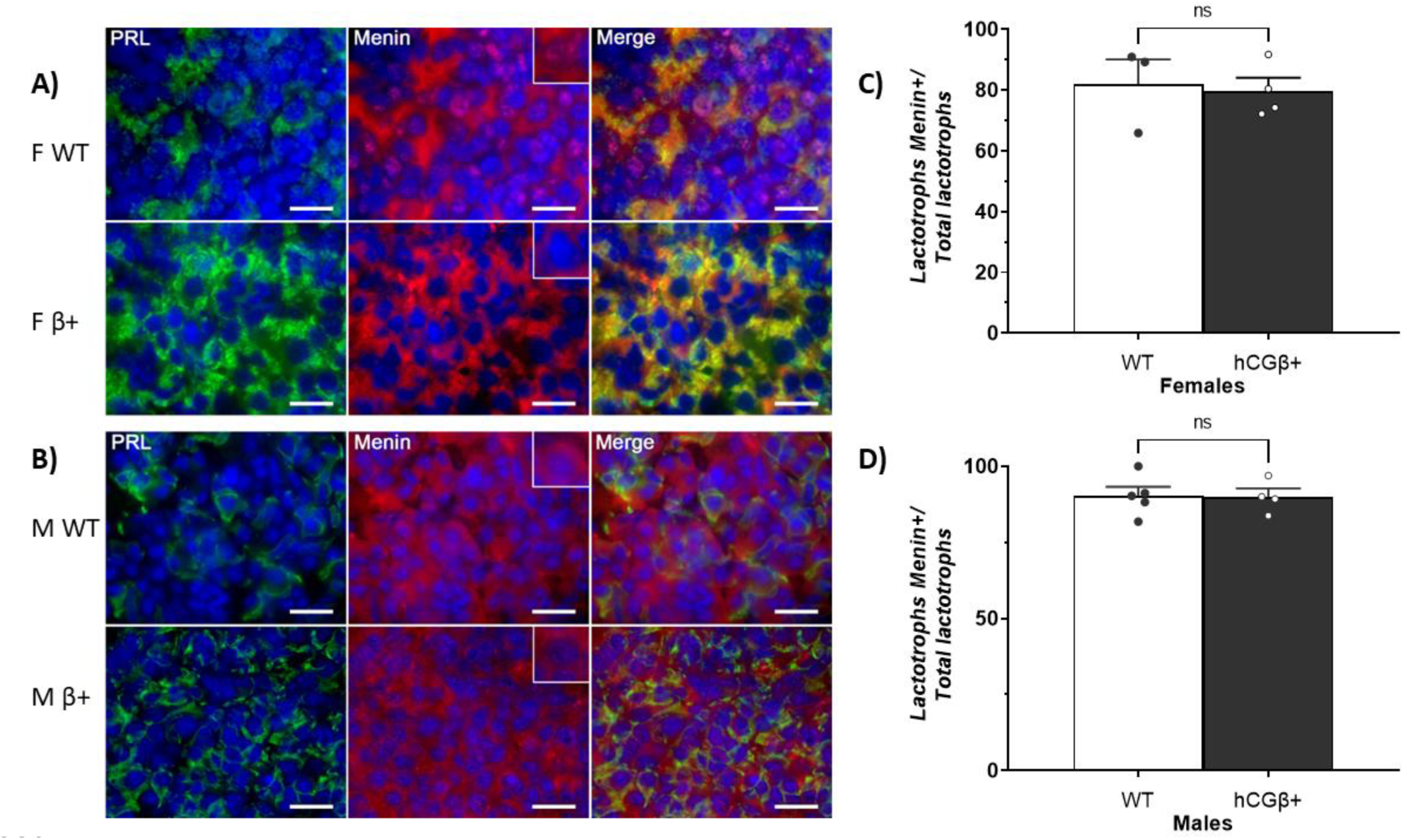
Menin protein expression in lactotrophs from hCGβ mice pituitaries. (A-B) Immunodetection of menin (red) and prolactin (green) by double indirect immunofluorescence in WT and hCGβ+ female (A) and male (B) mice pituitaries. Nuclei were immunostained with DAPI (blue). Scale bar = 20 μm. **(C-D)** Percentage of lactotrophs expressing Menin/total lactotrophs in female (C) and male (D) mice pituitaries. ns = not significant. Data analysed by Student t test. Data are expressed as mean ± SEM. n = 3-5/group.

However, we observed a pronounced difference in the subcellular localization of menin. In lactotrophs from WT female pituitaries of both animal models (Fig. 2A and 3A) and in lactotrophs from male pituitaries of both models (Fig. 2B and 3B), menin was localized in the cytoplasm and nucleus. In contrast, in lactotrophs from prolactinomas (Drd2KO and hCGβ⁺ females), menin was detected exclusively in the cytoplasm with an evident loss of nuclear localization.

### Dopamine and estradiol control MEN1 subcellular localization in a sex-specific manner

Upon observing alterations in menin subcellular localization, we next sought to identify factors involved in regulating its intracellular distribution. Given that dopamine and estradiol are key regulators of lactotroph function, we conducted *in vivo* experiments in Drd2 WT mice, both sexes (Figure 4).

**Figure 4:**
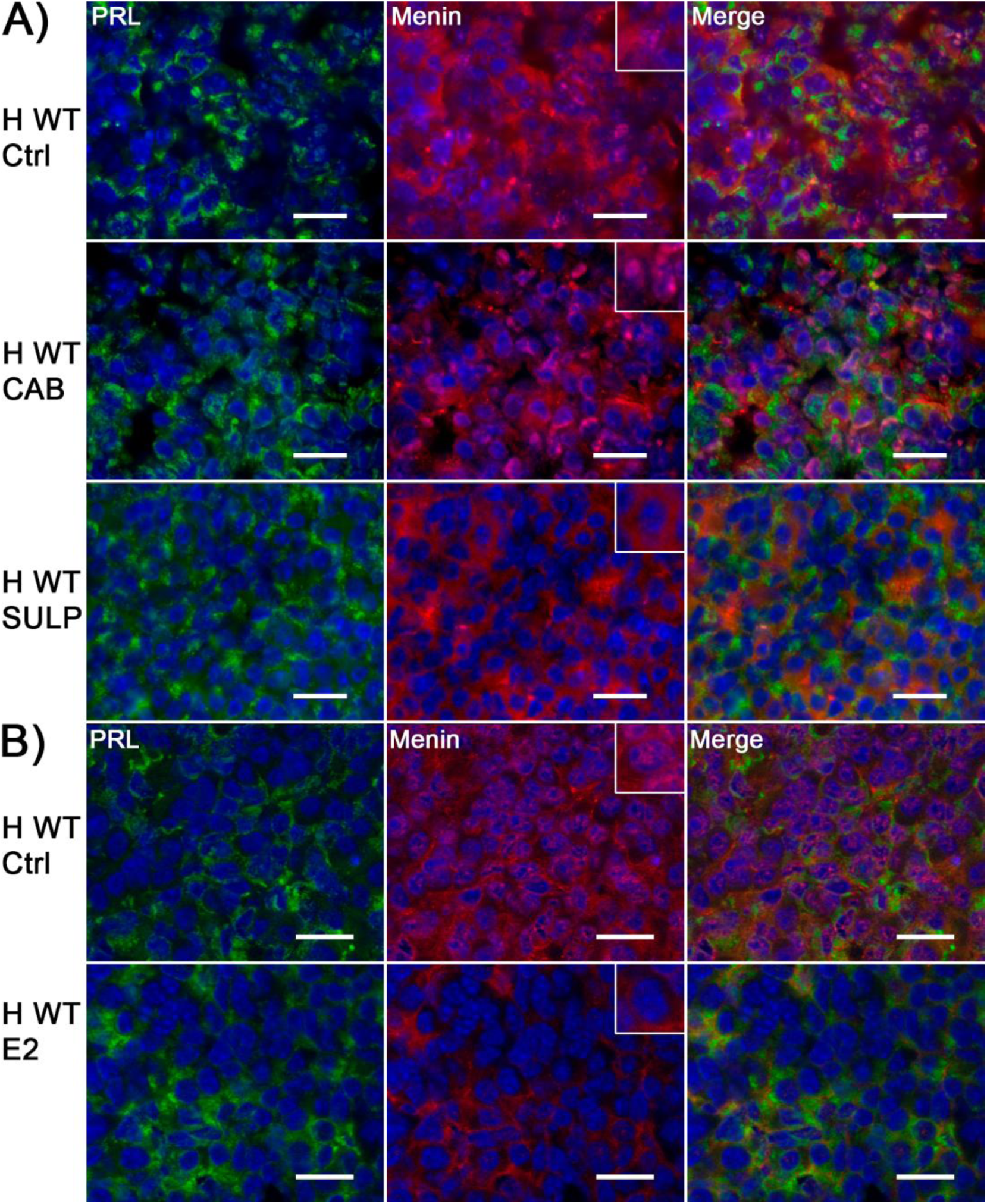
Dopamine and estradiol effect on menin subcellular localisation in WT female mice pituitaries (Drd2 model). Immunodetection of menin (red) and prolactin (green) by double indirect immunofluorescence in WT female mice pituitaries. Nuclei were immunostained with DAPI (blue). Scale bar = 20 μm. **(A)** *In vivo* treatment with cabergoline (CAB) or sulpiride (SULP). **(B)** *In vivo* treatment with estradiol (E2). Ctrl: control groups. n = 3-4/group.

As shown in figure 4A, cabergoline treatment preserved the nuclear localization of menin in WT female lactotrophs. In contrast, treatment with sulpiride or estradiol (Fig. 4B) led to the loss of nuclear menin expression. These results suggest that dopamine promotes, whereas estradiol inhibits, the nuclear localization of menin in female lactotrophs.

On the other hand, treatment with either cabergoline or sulpiride in WT males (Figure 5) did not alter the subcellular localization of menin in lactotrophs, suggesting that menin localization is not regulated by dopamine in this sex. However, estradiol administration induces only a partial loss of nuclear menin in lactotrophs from this sex.

**Figure 5:**
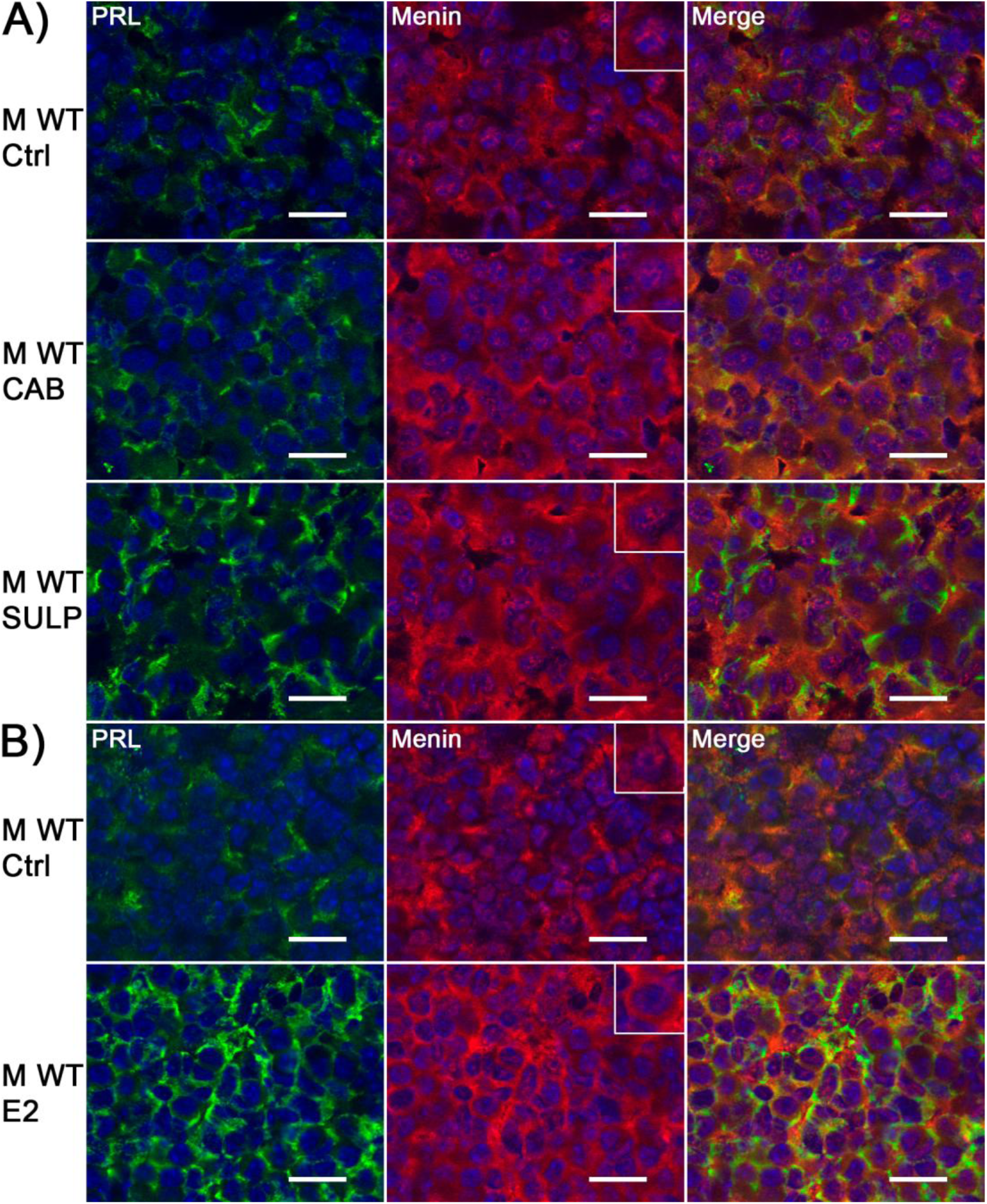
Dopamine and estradiol effect on menin subcellular localisation in WT male mice pituitaries (Drd2 model). Immunodetection of menin (red) and prolactin (green) by double indirect immunofluorescence in WT male mice pituitaries. Nuclei were immunostained with DAPI (blue). Scale bar = 20 μm. **(A)** *In vivo* treatment with cabergoline (CAB) or sulpiride (SULP). **(B)** *In vivo* treatment with estradiol (E2). Ctrl: control groups n = 3-4/group.

To further investigate the role of serum estradiol levels in modulating menin subcellular localization, we performed bilateral ovariectomy (OVX) in female hCGβ⁺ mice at two months of age, and pituitaries were studied at 6 months of age. The chronic absence of ovarian estradiol avoids prolactinoma development (22). As shown in figure 6, in those OVX-transgenic females pituitary MEN1 maintains the cytoplasmic but also the nuclear localization in lactotrophs, further supporting the importance of circulating estradiol in regulating menin subcellular distribution. Overall, these findings indicate that dopamine and estradiol regulate Men1 subcellular localization in lactotrophs, and, moreover, this effect is sex-dependent.

**Figure 6:**
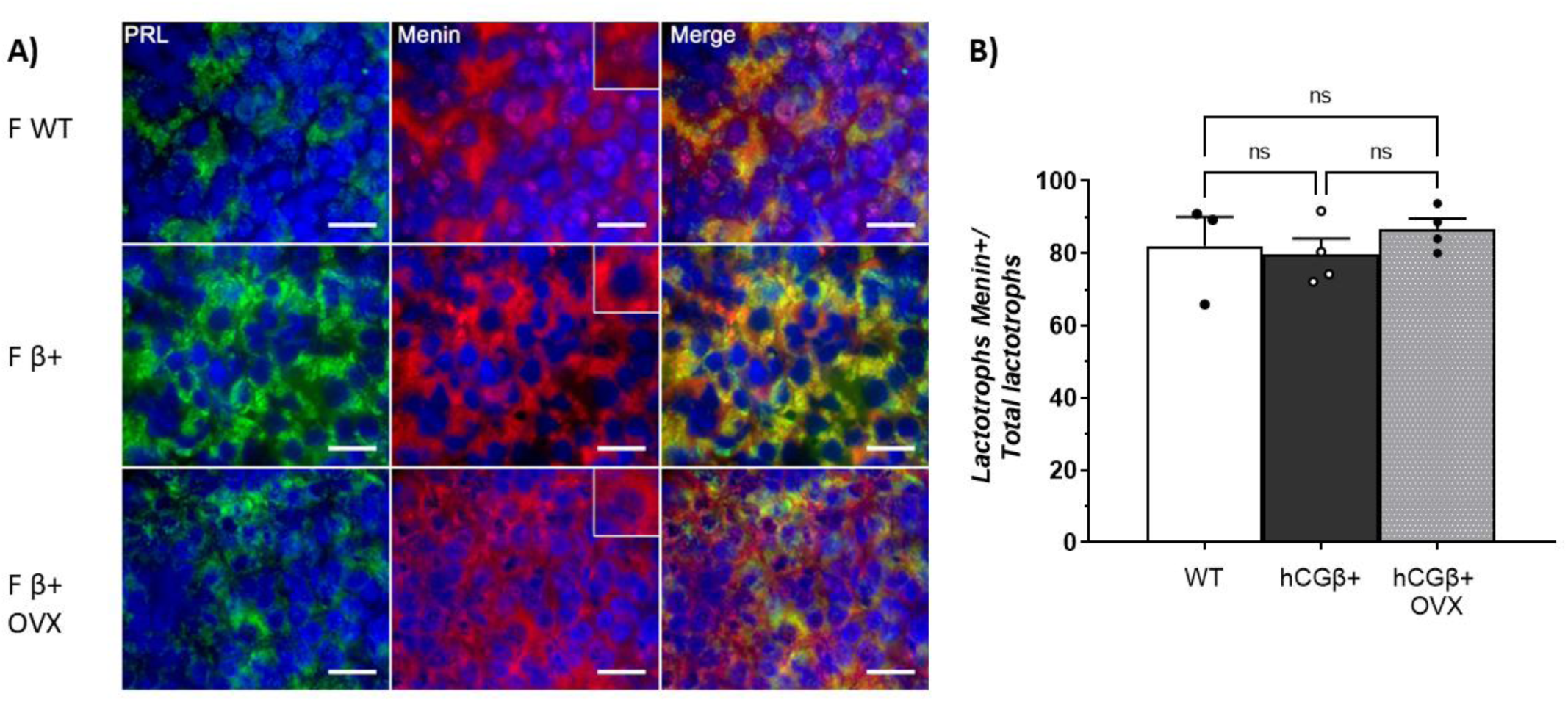
Effect of prepuberal OVX on menin subcellular localisation in hCGβ+ female mice pituitaries. **(A)** Immunodetection of menin (red) and prolactin (green) by double indirect immunofluorescence in WT, hCGβ+ and hCGβ+OVX mice pituitaries. Nuclei were immunostained with DAPI (blue). Scale bar = 20 μm. **(B)** Percentage of lactotrophs expressing Menin/total lactotrophs in WT, hCGβ+ and hCGβ+OVX mice pituitaries. ns = not significant. Data analysed by one-way ANOVA. Data are expressed as mean ± SEM. n = 3-4/group.

### Alterations in menin downstream effectors

The loss of nuclear menin expression in lactotrophs from transgenic female mice suggests a dysregulation of menin’s downstream effectors, which control cell proliferation and could be involved in prolactinoma development.

To further evaluate the impact of the loss of nuclear menin expression in prolactinomas, we analyzed several menin downstream effectors in the experimental groups.

### **a-** *Cdkn1B* gene expression and p27 protein expression

When we quantified pituitary *Cdkn1B* gene expression (Figure 7), one of the menin target genes encoding the CDK Inhibitor (p27 protein), we observed no differences among genotypes in female pituitaries from the Drd2 model. However, we found a decrease in *Cdkn1b* levels in hCGβ+ female pituitaries compared with their respective WTs. On the other hand, we did not observe differences in pituitary *Cdkn1b* expression among genotypes in males from the two animal models.

**Figure 7:**
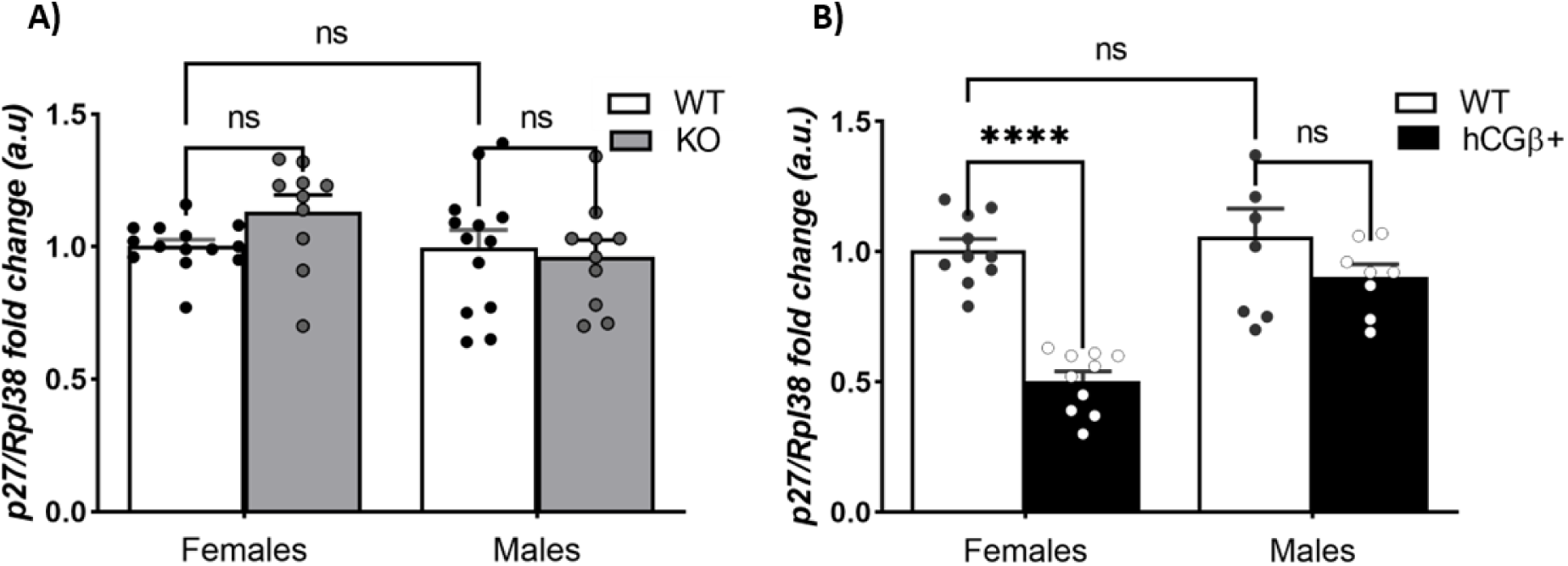
Pituitary mRNA expression of CDKN1B *(p27^Kip1^)* in Drd2 and hCGβ mouse models. mRNA transcripts were amplified with specific primers by qRT-PCR and normalized to *Rpl38*. Results are expressed relative to those for WT females. **(A)** 8-month-old Drd2 mice. Interaction ns. **(B)** 6-month-old hCGβ mice. Interaction *p* < 0,01. \*\*\*\**p* < 0,0001 hCGβ+-females vs WT-females. ns = not significant. Data analysed by two-way ANOVA (sex x genotype), followed by a Tuckey’s *post hoc* analysis when the interaction effect was significant. Data are expressed as mean ± SEM. n=7-13/group.

However, when we evaluated and quantified p27 protein expression by double immunofluorescence with PRL (Figure 8), we observed a decrease in the percentage of lactotrophs (PRL+) expressing p27 in prolactinomas from Drd2KO females compared with WT pituitaries (Fig. 8A and C). Nevertheless, we found no significant changes in the percentage of PRL+/p27+ cells when comparing hCGβ+ females with their WT counterparts (Figure 9), even though a decrease in p27+ lactotrophs was observed in four of the six prolactinomas evaluated (Fig. 9A and C). In male pituitaries from both mouse models the proportion of PRL+/p27+ cells was lower than the observed in the respective female pituitaries, but with no genotype alterations (Fig. 8B and D; Fig. 9B and D).

**Figure 8:**
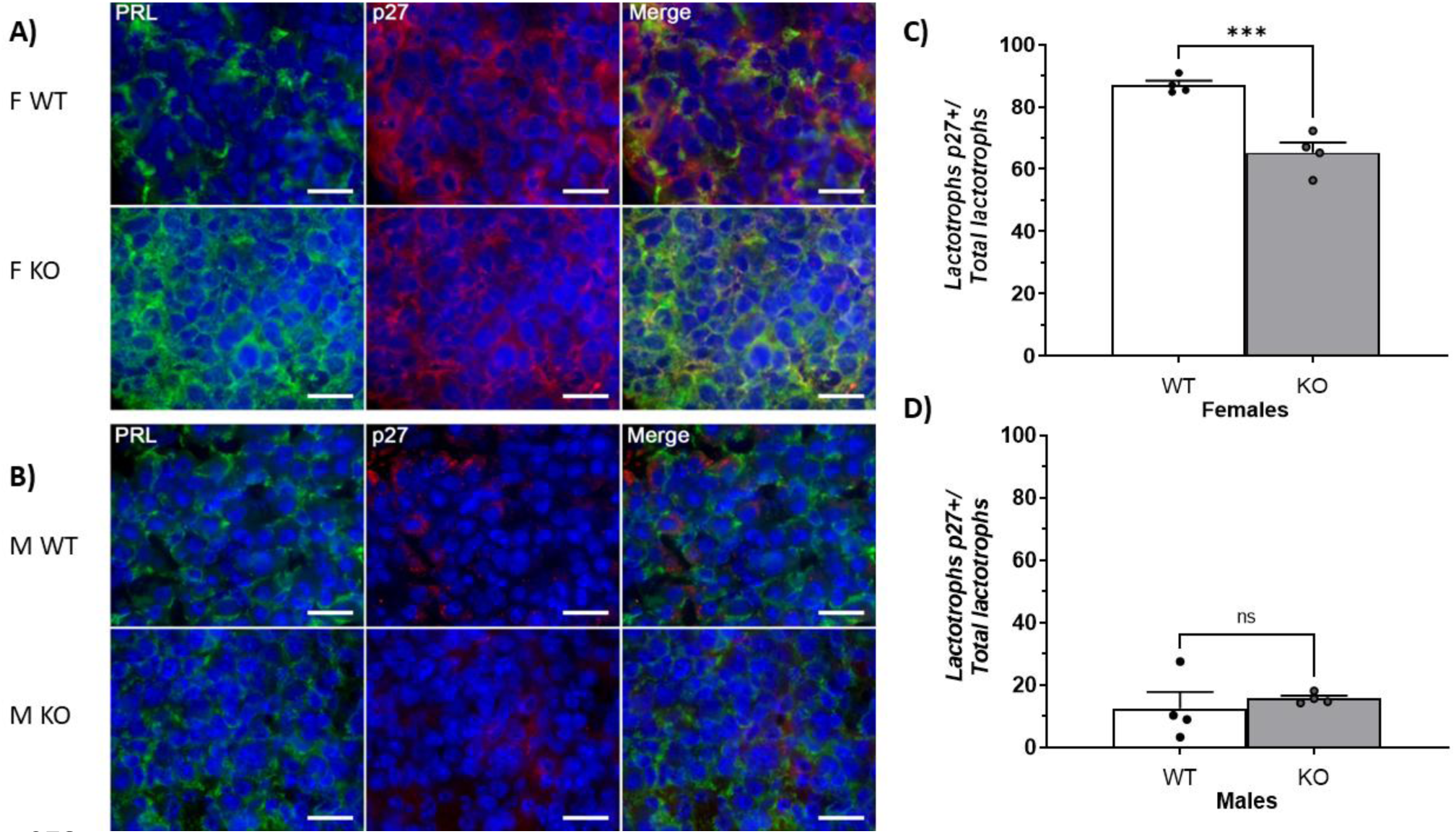
p27 protein expression in lactotrophs from Drd2 mice pituitaries. (A-B) Immunodetection of p27 (red) and prolactin (green) by double indirect immunofluorescence in WT and KO female (A), and male (B) mice pituitaries. Nuclei were immunostained with DAPI (blue). Scale bar = 20 μm. **(C-D)** Percentage of lactotrophs expressing p27/total lactotrophs in female (C) and male (D) mice pituitaries. \*\*\**p* < 0.001 KO-females vs WT-females. ns = not significant. Data analysed by Student t test. Data are expressed as mean ± SEM. n = 4/group.

**Figure 9:**
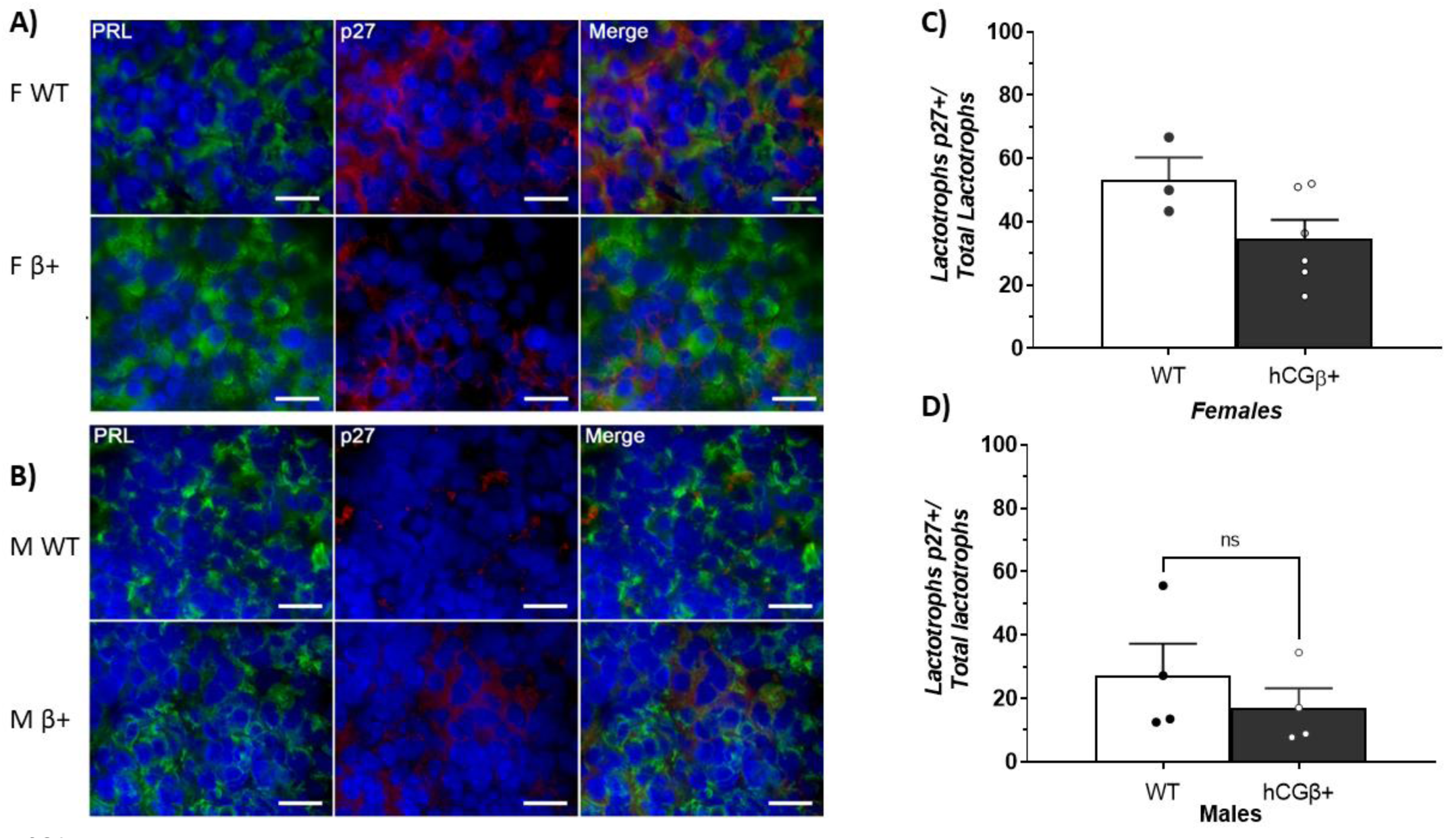
p27 protein expression in lactotrophs from hCGβ mice pituitaries. (A-B) Immunodetection of p27 (red) and prolactin (green) by double indirect immunofluorescence in WT and hCGβ+ female (A) and male (B) mice pituitaries. Nuclei were immunostained with DAPI (blue). Scale bar = 20 μm. **(C-D)** Percentage of lactotrophs expressing p27/total lactotrophs in female (C) and male (D) mice pituitaries. ns = not significant. Data analysed by Student t test. Data are expressed as mean ± SEM. n = 3-6/group.

### b-Cyclins*: Ccnd1* mRNA expression (Cyclin D1) and *Ccne1* mRNA expression(cyclin E1)

Next, we evaluated the pituitary mRNA expression of *Ccnd1* (Cyclin D1 gene) and *Ccne1* (Cyclin E1 gene) (Figure 10). We found increased *Ccnd1* levels in prolactinomas from the two mouse models (transgenic females) when compared with their WT counterparts (Fig. 10A and B), but no genotype differences in males. In addition, we identified higher levels of *Ccnd1* expression in male pituitaries from hCGβ model compared with their corresponding WT females. However, these sex differences were not observed in the Drd2 model.

**Figure 10:**
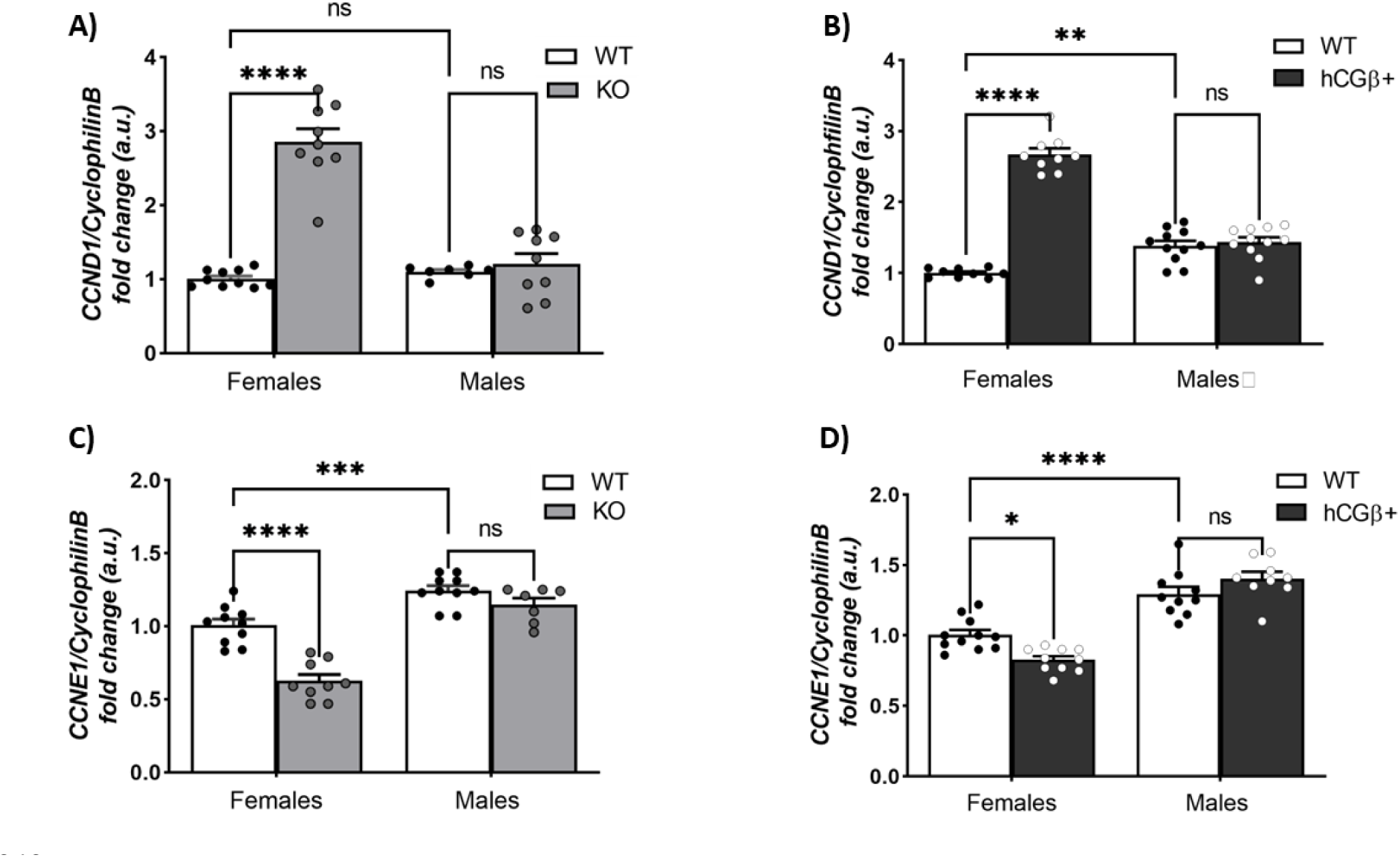
Pituitary mRNA expression of *CCND1 and CCNE1* in Drd2 and hCGβ mice. mRNA transcripts were amplified with specific primers by qRT-PCR and normalized to *CyclophilinB*. Results are expressed relative to those for WT females. **(A-B)** *CCND1* mRNA relative expression in 8-month-old Drd2 mice (A) and 6-month-old hCGβ mice (B). **(A)** Interaction *p* < 0,0001. \*\*\*\**p* < 0,0001 KO-females vs WT-females. **(B)** Interaction *p* < 0,0001. \*\*\*\**p* < 0,0001 hCGβ+-females vs WT-females; \*\**p* < 0,01 WT-males vs WT-females (sex difference). **(C-D)** *CCNE1* mRNA relative expression in 8-month-old Drd2 mice (C) and 6-month-old hCGβ+ mice (D). **(C)** Interaction *p* = 0,0016. \*\*\*\**p* < 0,0001 KO-females vs WT-females; \*\*\**p* < 0,001 WT-males vs WT-females (sex difference). **(D)** Interaction *p* = 0,0012. \**p* < 0,05 hCGβ+-females vs WT-females; \*\*\*\**p* < 0,0001 WT-males vs WT-females (sex difference). ns = not significant. Data analysed by two-way ANOVA (sex x genotype), followed by a Tuckey’s *post hoc* analysis when the interaction effect was significant. Data are expressed as mean ± SEM. n=7-10/group.

Regarding pituitary *Ccne1* expression, unexpectedly, we found decreased levels in Drd2KO and hCGβ+ female pituitaries (prolactinomas) compared with their WT counterparts (Fig. 10C and D). On the other hand, higher levels of pituitary *Ccne1* were observed in male pituitaries compared to their respective WT females in both mouse models, with no genotype differences.

### c-pAKT is overexpressed in prolactinomas

As mentioned, menin can interact and interfere with AKT phosphorylation status, a key factor in regulating cell proliferation (8). By double immunofluorescence, we found very low levels of pAKT (Ser473) in WT pituitaries, both models, both sexes, and restricted to the cytoplasm (Figure 11, B and D; Figure 12, B and D). However, we observed considerably high pAKT expression in lactotrophs from prolactinomas (transgenic female pituitaries, both models) compared to WTs (Fig. 11A and C; Fig. 12A and C), in both, the cytoplasm and nuclei, in agreement with the decreased levels of menin in tumoral pituitaries.

**Figure 11:**
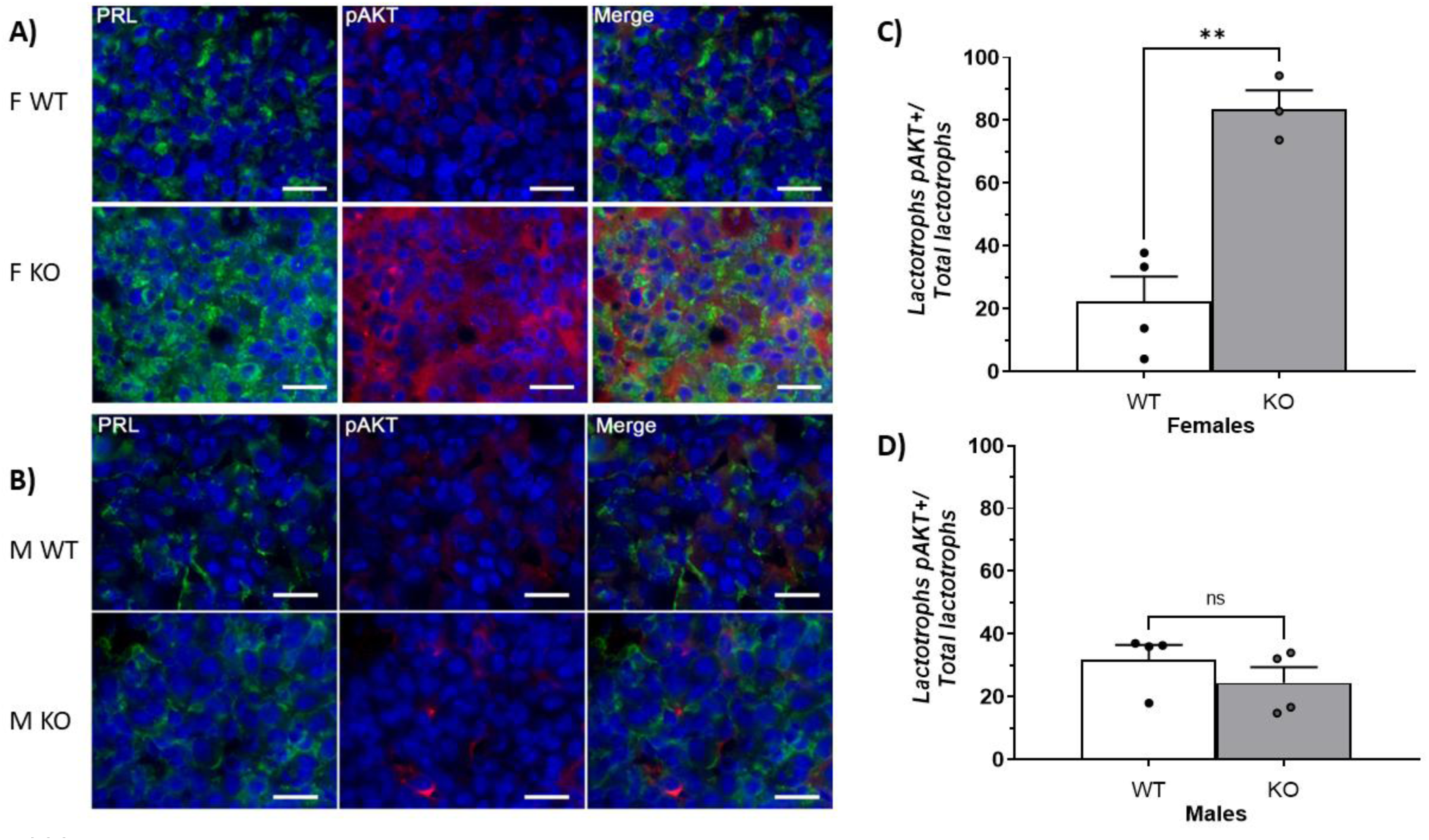
pAKT protein expression in lactotrophs from Drd2 mice pituitaries. (A-B) Immunodetection of pAKT (red) and prolactin (green) by double indirect immunofluorescence in WT and KO female (A) and male (B) mice pituitaries. Nuclei were immunostained with DAPI (blue). Scale bar = 20 μm. **(C-D)** Percentage of lactotrophs expressing pAKT/total lactotrophs in female (C) and male (D) mice pituitaries. \*\**p* < 0.01 KO-females vs WT-females. ns = not significant. Data analysed by Student t test. Data are expressed as mean ± SEM. n = 3-4/group.

**Figure 12:**
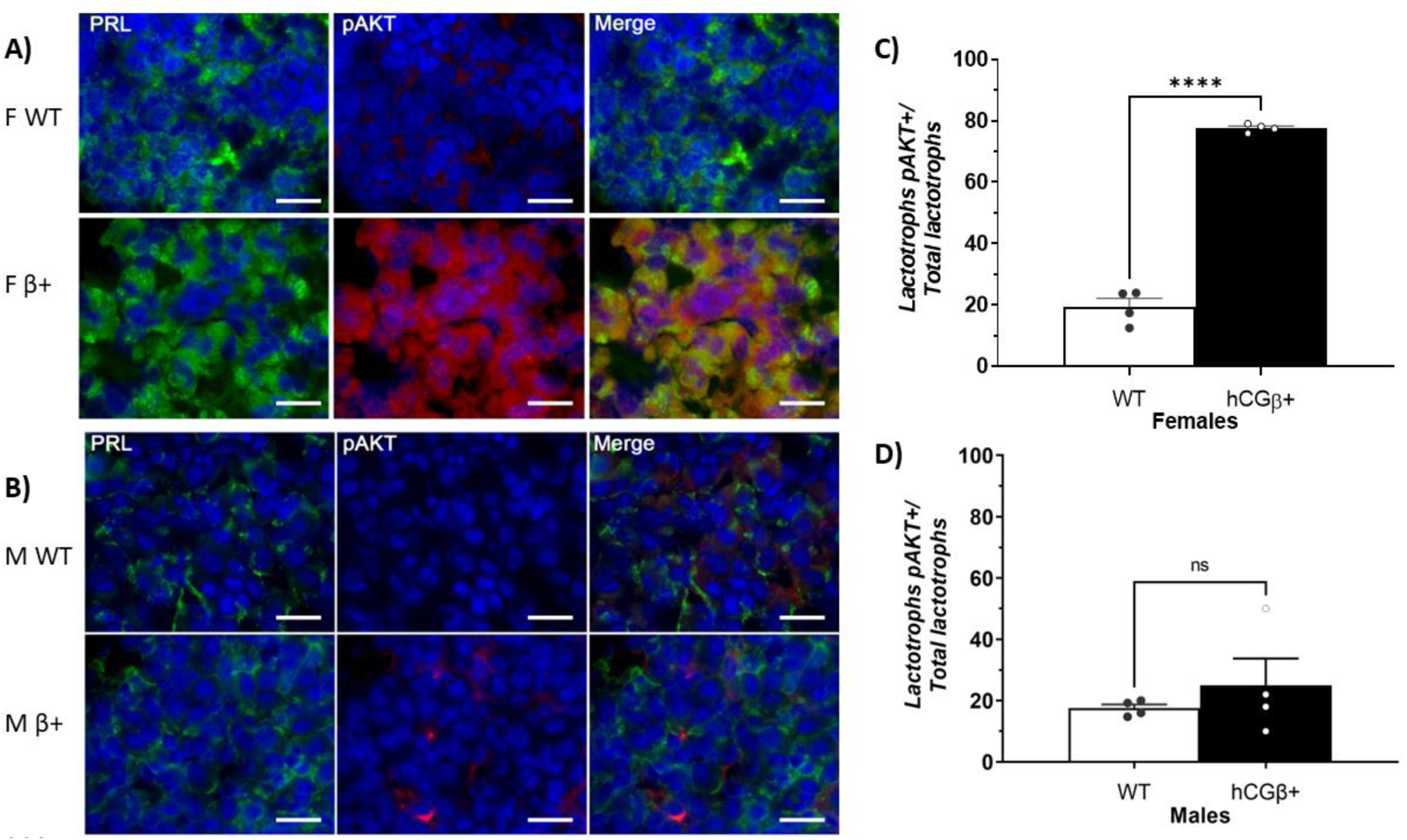
pAKT protein expression in lactotrophs from hCGβ mice pituitaries. (A-B) Immunodetection of pAKT (red) and prolactin (green) by double indirect immunofluorescence in WT and hCGβ+ female (A) and male (B) mice pituitaries. Nuclei were immunostained with DAPI (blue). Scale bar = 20 μm. **(C-D)** Percentage of lactotrophs expressing pAKT/total lactotrophs in female (C) and male (D) mice pituitaries. \*\*\*\**p* < 0.0001 hCGβ+-females vs WT-females. ns = not significant. Data analysed by Student t test. Data are expressed as mean ± SEM. n = 4/group.

Overall, these findings indicate that the loss on nuclear menin expression in tumoral lactotrophs disrupt menin regulatory pathways, and could be helping the tumorigenic process. Once again, those processes are sex-dependent.

### PTEN gene expression is decreased in prolactinomas from both mouse models

One of the best-characterized inhibitors of AKT phosphorylation is PTEN (23)(24), which cooperates with menin in suppressing lactotroph proliferation. When we analyzed pituitary *Pten* expression, we found significantly decreased levels in Drd2KO compared with WT‘s, with no effect of sex or interaction (Figure 13). In hCGβ mice, however, pituitary *Pten* expression was significantly higher in males than in females. This marked sex difference may have masked a genotype effect in females in the two-way ANOVA. Indeed, when females were analyzed separately using Student’s t-test, *Pten* expression was significantly decreased in hCGβ+ pituitaries compared with WT‘s.

**Figure 13:**
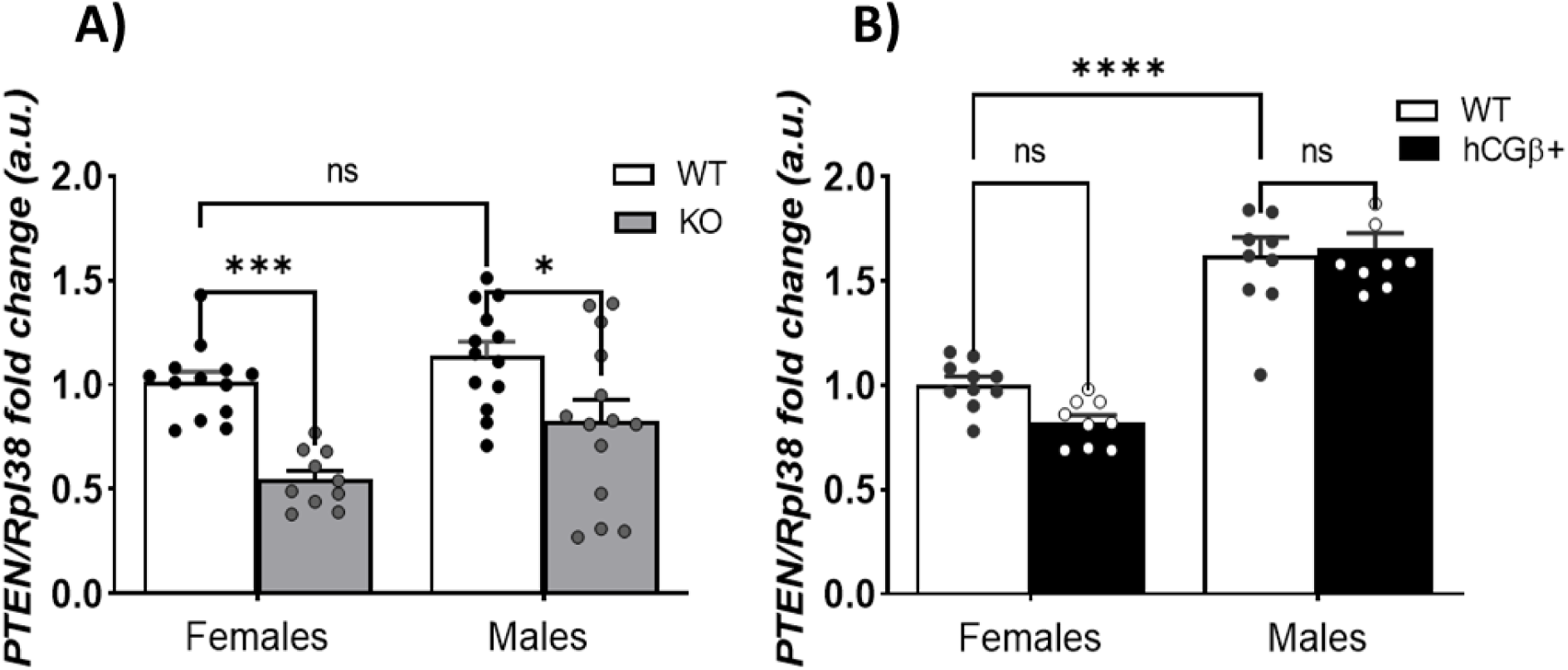
Pituitary mRNA expression of PTEN in Drd2 and hCGβ mice. mRNA transcripts were amplified with specific primers by qRT-PCR and normalized to *Rpl38*. Results are expressed relative to those for WT females. **(A)** 8-month-old Drd2 mice. Interaction ns. \*\*\**p* < 0,001 KO-females vs WT-females; \**p* < 0,05 KO-males vs WT-males. **(B)** 6-month-old hCGβ mice. Interaction ns. \*\*\*\**p* < 0,0001 WT-males vs WT-females (sex difference). (Insert: \*\**p* < 0,01 differences between hCGβ+-females vs WT-females by t test). ns = not significant. Data analysed by two-way ANOVA (sex x genotype). Data are expressed as mean ± SEM. n=9-14/group.

### Exploratory assessment of menin subcellular localization in human normal pituitaries and prolactinomas

To explore whether the alterations in menin subcellular localization observed in the murine models might also be observed in human prolactinomas, we analyzed MEN1 immunoreactivity in an available, but limited, set of human pituitary biopsies by immunofluorescence. In normal anterior pituitaries from both women and men, lactotrophs exhibited intense cytoplasmic prolactin immunoreactivity, consistent with the abundant mature secretory granules that characterize a distinctive feature of normal lactotrophs. On the other hand, menin displayed strong immunoreactivity detected in both, the nucleus and cytoplasm (Figure 14B). A similar distribution of PRL and menin was observed in prolactinomas from a male, and from a woman (Fig. 14A), both who were pre-treated with cabergoline before surgery. In contrast, a prolactinoma from a woman who had not received dopamine agonist therapy showed lactotrophs exhibiting a weak cytoplasmic prolactin immunostaining reflecting a depletion of prolactin-containing secretory granules due to an increased exocytotic activity and sustained hormone secretion as previously documented (25)(26). Interestingly, a noticeable loss of nuclear menin was observed, with the protein remaining disaggregated and predominantly in the cytoplasm. Although we acknowledge that the number of human samples is limited, these observations provide proof of concept in human prolactinomas because they closely mirror the findings obtained in the experimental mouse models. Specifically, strong nuclear menin expression was observed in normal pituitaries and treated prolactinomas from both sexes, whereas nuclear exclusion of menin was detected in the untreated prolactinoma from a female patient.

**Figure 14:**
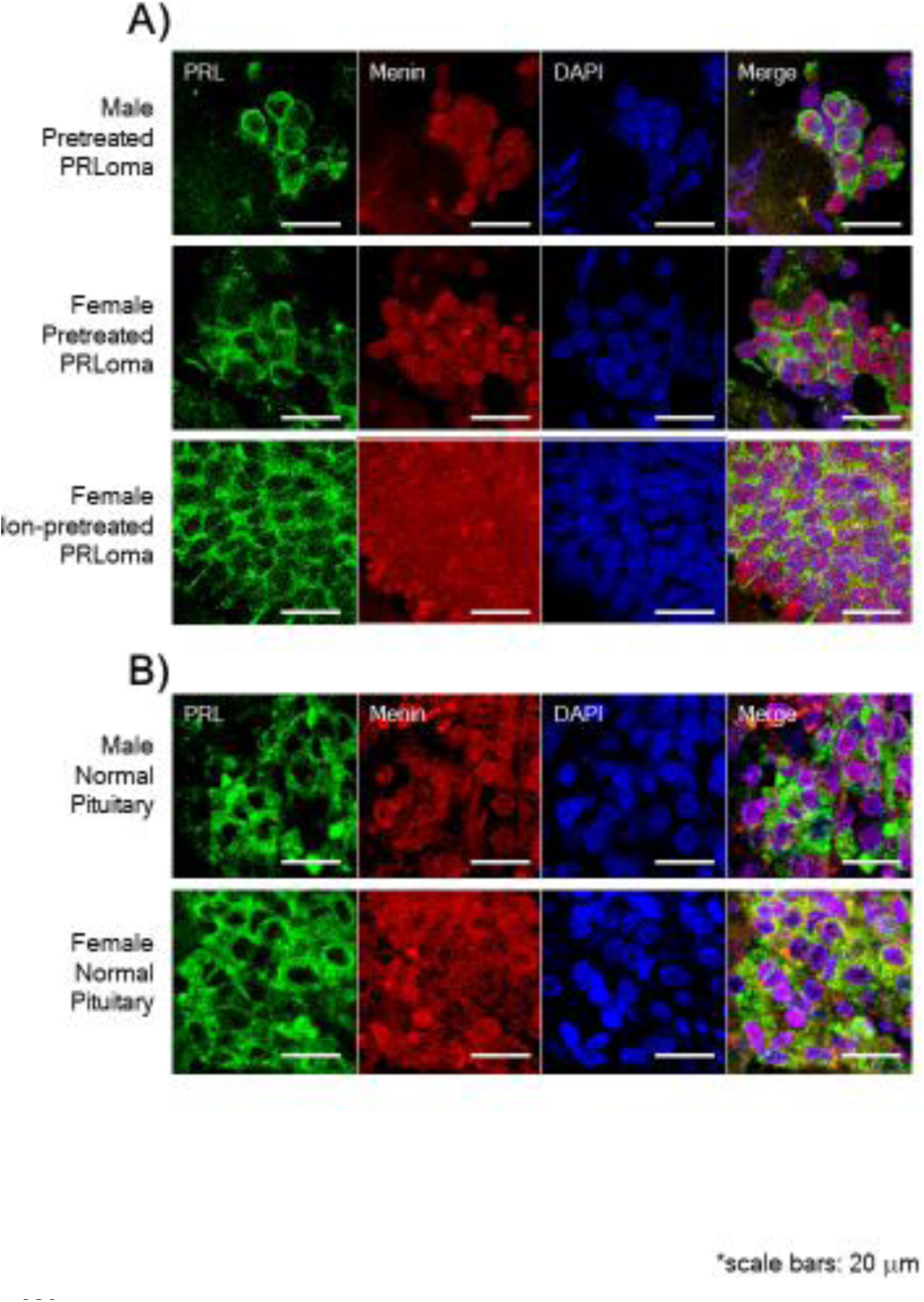
MEN1 expression in human biopsies from prolactinomas (A) and normal pituitaries (B): Immunodetection of menin (red) and prolactin (green) by double indirect immunofluorescence in human pituitaries. Nuclei were immunostained with DAPI (blue). Scale bar = 20 μm

## DISCUSSION

It is well-known that the germline loss-of-function mutations in *Men1* cause Multiple Endocrine Neoplasia type 1 (MEN1) syndrome. This condition predisposes MEN1 individuals to neuroendocrine tumours, including prolactinomas (2,27). Prolactinomas represent the most prevalent type of secreting pituitary tumours, comprising around 40% of all of them, and are characterized by lactotroph hyperplasia and excessive prolactin secretion (28). However, the role of menin in the pathophysiology of sporadic prolactinomas, which occur in the absence of *Men1* mutations, remains less clear.

A major finding of the present work is that prolactinoma development, in the mouse models studied, occurred in the absence of detectable changes in *Men1* expression. Instead, tumoral lactotrophs exhibited a striking loss of nuclear menin localization while retaining cytoplasmic expression. This observation suggests that menin dysfunction in sporadic prolactinomas does not result from genetic loss of *Men1* but rather from altered intracellular trafficking and subcellular compartmentalization.

The biological consequences of this phenomenon are potentially equivalent to MEN1 loss-of-function. Menin exerts many of its tumour suppressor actions within the nucleus, wherein it regulates transcriptional programs controlling cell cycle progression, differentiation, and growth suppression (3–7). Therefore, exclusion of menin from the nucleus may generate a state of functional impairment of menin-dependent nuclear signalling despite preservation of MEN1 gene expression.

This concept provides a mechanistic framework linking hereditary and sporadic prolactinomas. While MEN1-associated tumours lose menin activity through genetic alterations, sporadic prolactinomas may achieve a similar biological outcome through disruption of menin nuclear localization.

Then, the central paradox was to find normal *Men1* mRNA levels in prolactinomas but with loss of function in nucleus. This is a critical point because it shifts the focus of the pathology in these models (prolactinoma development) away from gene expression (the classic mechanism in MEN1 syndrome) to post-translational regulation and protein function involved in our experimental mouse models. In the present work we highlight the importance of observing alterations in the menin subcellular localization and the impact on its functions. Even though the protein is present in lactotrophs (in the cytoplasm), it is not in the nucleus, wherein menin performs its primary function as a transcriptional regulator of its nuclear targets. This explains how tumour suppression fails even with normal *Men1* gene expression.

Menin is known to regulate cell cycle progression through the transcriptional activation of CDKIs, such as p27 (*Cdkn1b*), or *Ccnd1*, a gene that encodes Cyclin D1 crucial for cell cycle progression (5–7). In hCGβ+ tumoral pituitaries, our results confirmed these mechanisms, showing decreased p27 gene expression together with increased *Ccnd1*, consistent with the classic consequence of impaired nuclear menin function. In contrast, in Drd2KO pituitaries, p27 gene expression remained unchanged despite the loss of nuclear menin. However, we observed reduced p27 protein levels in this model, which may instead be explained by the strong activation of the AKT pathway, leading to decreased p27 protein stability (29). In both models, the concomitant increase in *Ccnd1* highlights a powerful driver of cell cycle progression.

The effect observed on the PTEN/pAKT pathway is another crucial evidence that could be involved in the loss of several menin functions in prolactinomas. Menin-AKT interaction is well documented in the cytoplasm, wherein menin regulates AKT function. Menin acts as a scaffold protein to bind and sequester inactive AKT, avoiding its translocation from the cytoplasm to the plasma membrane and preventing AKT activation (8). Additionally, menin cooperates with the tumour suppressor PTEN to limit the mitogenic action of pAKT, a potent pro-survival and pro-proliferative signal (24). The decrease in *PTEN* expression observed in prolactinomas was concomitant with pAKT overexpression and its localization to both the cytoplasm and nucleus, thereby promoting a potent proliferative signal in tumoral pituitaries.

Beyond its effects on the p27 and PTEN/AKT pathways, the loss of nuclear menin in lactotrophs from prolactinomas offers a compelling molecular explanation for the dysfunction of other key tumour-suppressive systems. Previous works reported impaired activin and TGFβ signalling in female prolactinomas (30–32). Menin is an essential component of SMAD-dependent transcriptional complexes in lactotrophs. TGFβ1 and activins induce SMAD2/3 phosphorylation, leading to their association with SMAD4 in the cytoplasm. Menin physically interacts with the activated SMAD complex and is required for its nuclear translocation and transcriptional activity on TGFβ/activin target genes. Moreover, loss of menin impairs SMAD3 DNA binding and disrupts the antiproliferative TGFβ/activin signalling pathway (33)(34). Consequently, loss of nuclear menin provides a direct mechanistic explanation for the reduced responsiveness of tumoral lactotrophs to activins and TGFβ1 previously described in animal models of prolactinoma. Thus, the nuclear exclusion of menin may represent a common upstream event integrating alterations in cell cycle regulation, AKT activation, and TGFβ/activin tumour suppressor signalling. Consistently, we demonstrate here that, in male pituitaries, menin localizes to both the cytoplasm and nucleus regardless of genotype, helping to maintain those inhibitory signals, protecting males from pituitary tumorigenesis (32,35,36). All these data reinforce the notion that, in our models and at the ages studied, males possess more robust intra-pituitary inhibitory mechanisms that counteract prolactinoma development.

Additionally, our present results identify, for the first time, that menin subcellular localization is dynamically regulated by endocrine signals: dopamine and estradiol control menin intracellular localization in lactotrophs, expanding on previous findings. Chronic estradiol treatment is well known to induce prolactinoma development (37). It was previously described that estradiol impairs the inhibitory action of activins and TGFβ1 (31,36,38,39). We now demonstrate that estradiol induces the nuclear exclusion of menin, leading to alterations in key cell cycle effectors. This mechanism may also contribute to the previously described impairment of inhibitory signalling pathways, including the TGF-β/activin–pSMAD2/3 axis (35,40). The prevention of all these alterations by prepuberal ovariectomy in transgenic females further confirms the central role of estradiol in those processes (41). Conversely, dopamine preserves nuclear menin in female pituitaries, and blocking dopamine action with sulpiride induces the loss of nuclear menin localization. Once again, these findings are consistent with previous studies demonstrating a positive effect of dopamine on TGFβ1 biological activity in lactotroph cells (38)(42), and provides a novel additional knowledge on the molecular mechanism for dopamine’s tumour-suppressive effects.

Another noteworthy observation was the absence of any effect of dopamine on menin subcellular localization in male pituitaries. Sex differences in estradiol effects and dopaminergic tone at the pituitary level have long been recognized (43), with basal hypothalamic dopamine activity and lactotroph responsiveness approximately five times higher in females than in males. Consistent with this, Drd2 disruption exerts a more profound impact on pituitary function in female mice than in males (44)(45).

A particularly relevant finding of the present study is that the alterations in menin subcellular localization identified in two independent mouse models were also observed in human prolactinomas. Specifically, we found that menin displayed both nuclear and cytoplasmic localization in normal human pituitaries of both genders. Moreover, menin was also observed with nuclear localization in prolactinomas from dopamine agonist-treated patients, consistent with our experimental evidence showing that dopamine preserves nuclear menin localization. Interestingly, we observed a partial and scattered loss of nuclear menin expression in an untreated female prolactinoma. Although these observations derive from a limited number of surgical biopsies, they provide an important proof of concept/translational validation of our experimental findings in humans and suggest that loss of nuclear menin is not merely a feature of animal models but may represent a clinically relevant mechanism in sporadic human prolactinomas. Furthermore, the association between dopamine agonist treatment and preserved nuclear menin localization raises the possibility that maintenance of nuclear menin may contribute to the antitumoral effects of these drugs.

In conclusion, we provide compelling evidenced demonstrating that nuclear exclusion of menin is a previously unrecognized molecular mechanism associated with sporadic prolactinoma development. Unlike MEN1-associated tumours, prolactinomas from two independent non-MEN1 mouse models retain MEN1 (gene and protein) expression while exhibiting loss of nuclear menin localization, generating a state of functional impairment of menin-dependent nuclear signalling. This alteration is associated with reduced p27 and PTEN signalling, increased AKT activation, enhanced cyclin D1 expression, and impaired activin/TGFβ tumour suppressor pathways. The ability of dopamine agonism and ovariectomy to preserve or restore nuclear menin localization further identifies menin trafficking as a potential therapeutic target for prolactinomas, particularly those resistant to conventional dopaminergic therapies.

## All authors declare that there is no conflict of interest Funding

This work was supported by BIGAND Fundation, Buenos Aires Argentina, to GDT; Agencia Nacional de Promoción Científica y Técnica, Buenos Aires, Argentina (grant PICT 2021 N0110 to GDT); René Barón Fundation, Williams Fundation, Florencio Fiorini Fundation to GDT.

Junta de Andalucía (PI-0117-2025, DGP_PIDI_2024_00134, BIO-0139), FSEEN, ISCIII (CD24_00240) and CIBERobn to RML. CIBER is an initiative of ISC-III, “Ministerio de Sanidad, Servicios Sociales e Igualdad”, Spain.

## AI Disclosure Statement

The authors used artificial intelligence (AI) tools (CHAT GPT) to assist with English language editing and to support the preparation of the graphical abstract. All scientific content, data analysis, interpretation of results, and conclusions were developed and reviewed by the authors, who take full responsibility for the accuracy and integrity of the work.

## Acknowledgements

We deeply thank all the patients and their families for generously donating human samples. Special thanks to the staff of Biobank of Andalusian Public Health System Biobank [Reina Sofia University Hospital (Córdoba node)]

